# Corticostriatal oscillations predict high vs. low drinkers in a preclinical model of limited access alcohol consumption

**DOI:** 10.1101/293597

**Authors:** Angela M. Henricks, Lucas L. Dwiel, Nicholas H. Deveau, Amanda A. Simon, Metztli J. Ruiz-Jaquez, Alan I. Green, Wilder T. Doucette

## Abstract

Individuals differ in their vulnerability to develop alcohol dependence that are determined by innate and environmental factors. The corticostriatal circuit is heavily involved in the development of alcohol dependence and may contain neural information regarding vulnerability to drink excessively. In the current experiment, we hypothesized that we could characterize high and low alcohol-drinking rats (HD and LD, respectively) based on corticostriatal oscillations, and that these subgroups would differentially respond to corticostriatal brain stimulation. Rats were trained to drink 10% alcohol in a limited access paradigm. In separate sessions, local field potentials (LFPs) were recorded from the nucleus accumbens shell (NAcSh) and medial prefrontal cortex (mPFC) of male Sprague-Dawley rats (n=13). Based on training alcohol consumption levels, we classified rats using a median split as HD or LD. Then, using machine-learning, we built predictive models to classify rats as HD or LD by corticostriatal LFPs and compared the model performance from real data to the performance of models built on data permutations. Additionally, we explored the impact of NAcSh or mPFC stimulation on alcohol consumption in HD vs. LD. Corticostriatal LFPs were able predict HD vs. LD group classification with greater accuracy than expected by chance (>80% accuracy). Additionally, NAcSh stimulation significantly reduced alcohol consumption in HD, but not LD (p<0.05), while mPFC stimulation did not alter drinking behavior in either HD or LD (p>0.05). These data collectively show that the corticostriatal circuit is differentially involved in regulating alcohol intake in HD vs. LD rats, and suggests that corticostriatal activity may have the potential to predict a vulnerability to develop alcohol dependence in a clinical population.

## Introduction

Excessive alcohol consumption is a major health concern in the United States, leading to approximately 88,000 deaths per year (Centers for Disease Control and Prevention, 2013), but only a small proportion of individuals who drink alcohol become dependent later in adulthood (Costanzo et al., 2007). Both innate and environmental risk factors are associated with the development of alcohol dependence which manifest within the brain (Hägele, Friedel, Kienast, & Kiefer, 2014; Matošić, Marušić, Vidrih, Kovak-Mufić, & Cicin-Šain, 2016). Even within outbred rats, there are significant variations in alcohol consumption levels. Rats categorized as high or low alcohol drinkers (HD and LD, respectively) display differences in anxiety, impulsivity, and cognitive behaviors compared to low drinkers (Pratt, Gates, Smith, & Amit, 2002; Sharko, Kaigler, Fadel, & Wilson, 2013; Wilhelm & Mitchell, 2008), as well as differences in gene expression known to influence appetitive behavior (Morganstern, Liang, Ye, Karatayev, & Leibowitz, 2012). Importantly, HD rats show behavioral phenotypes that may be associated with an increased vulnerability to develop addiction (Spoelder et al., 2017). Additional work is needed, however, to understand how systems-level neural activity relates to the HD phenotype and the predictive behavioral phenotypes.

Previous research indicates that a history of alcohol use is associated with dysregulation in the corticostriatal circuit (Broadwater et al., 2018; Camchong, Stenger, & Fein, 2013), including the nucleus accumbens (NAc) and both human and rat medial prefrontal cortex (mPFC). The NAc integrates cortical inputs and indirectly sends feedback to the mPFC (Goto & Grace, 2008), and is particularly important in the motivating and rewarding properties of abused drugs (Koob & Volkow, 2010). The mPFC is activated in response to reward-related cues, and it has been suggested that deficits in the ability to inhibit responses to drug and associated cues arises from reduced top-down control of the mPFC to striatal regions (Goldstein & Volkow, 2002). Additionally, stimulation of the NAc has recently been used to reduce alcohol consumption in both humans and rodents (Henderson et al., 2010; Knapp, Tozier, Pak, Ciraulo, & Kornetsky, 2009; Voges, Müller, Bogerts, Münte, & Heinze, 2013), though it is important to note that brain stimulation suffers from the same highly variable treatment outcomes observed with other psychiatric treatments. However, our previous research with binge eating suggests neural oscillations from the NAc can provide systems-level information regarding individual variability in behavior (unpublished results). The corticostriatal circuit is therefore an important target for studies aimed at understanding how risk factors leading to alcohol dependence become instantiated in the brain.

In the current experiment, we hypothesized that we could predict which rats were high alcohol drinkers (HD) or low alcohol drinkers (LD) using local field potential (LFP) oscillations recorded within the corticostriatal network, and that these subgroups would respond differentially to cortical and striatal stimulation. We recorded LFPs from two corticostriatal brain regions (NAc shell [NAcSh] and rat mPFC [infralimbic cortex and prelimbic cortex]) and subsequently treated rats with high frequency brain stimulation in each of those regions (separately) during alcohol drinking sessions. We theorized that variation in the effect of stimulation on alcohol behavior in HD vs. LD may, in part, be related to individual differences in the networks that underpin alcohol consumption.

## Materials and methods

### Animals

Male Sprague-Dawley rats (n=13) were purchased from Harlan (South Easton, MA, USA) and arrived on postnatal day 60. All animals were housed individually on a reverse 12-hour light cycle with *ad libitum* access to food and water. All experiments were carried out in accordance with the National Institute of Health Guide for the Care and Use of Laboratory Animals (NIH Publications No. 80-23) and were approved by the Institutional Animal Care and Use Committee of Dartmouth College.

### Electrode implantation

Electrodes were designed and constructed in-house and were similar to those used in our previous publication (W T Doucette, Khokhar, & Green, 2015). Following one week of habituation to the animal facility, animals were anesthetized with isoflurane gas (4% induction, 2% maintenance) and mounted in a stereotaxic frame. Custom electrodes were implanted bilaterally targeting NAcSh (from bregma: DV-8mm; AP +1.2mm; ML +/-1.0mm) and mPFC (from bregma: DV-5mm; AP +3.7mm; ML +/-0.75mm). Four stainless steel skull screws were placed around the electrode site and dental cement (Dentsply, York, PA, USA) was applied to secure the electrodes in place. Animals were allowed to recover for at least one week prior to being tethered to the recording apparatus and trained to consume a 10% alcohol solution.

### Alcohol consumption training

Animals were trained to drink 10% alcohol in a limited access paradigm. Three days per week (M, W, F) animals were transferred from their home cage to custom stimulation chambers, tethered to stimulation cables, and given free access to 10% alcohol for 90 minutes. Animal weights and the volume of alcohol consumed was measured following each session in order to calculate g/kg of alcohol consumed. Animals were allowed to drink while plugged in to stimulation cables, without stimulation, for 12 sessions.

### Local field potential recordings

Prior to brain stimulation sessions (but after exposure to alcohol), animals were tethered for LFP recording in a chamber that was distinct from the alcohol consumption and stimulation chamber. Animals engaged in free behavior while tethered through a commutator to a Plexon data acquisition system and video was recorded that was time-synchronized to the LFP data (Plexon, Plano, Tx) during two, 30-minute sessions. Noise free data from the entire 30-minute recording session were analyzed using established frequency ranges from the rodent literature (listed below) and standard LFP signal processing was used to characterize the power spectral densities (PSDs) within, and coherence between brain regions (bilateral NAcSh and mPFC) for each animal using custom code written for Matlab R2015b.

A fourth order Chebychev type I notch filter centered at 60 Hz was applied to all of the data to account for 60 Hz line noise. The data was then down-sampled by a factor of five from 2 kHz to 400 Hz. A threshold of ± 2 mV was used to identify noise artifacts and remove data using intervals 12.5 milliseconds before and 40 seconds after the artifacts. To capture the power and coherence dynamics of the signal, we used only behavioral epochs that were at least 3 seconds long. For epochs that were longer than 3 seconds, we segmented them into 3-second sections removing the remainder to keep all of the data continuous over the same amount of time.

PSDs were computed using MATLAB’s *pwelch* function using a 1.6 second Hamming window with 50% overlap. The PSDs for each 3-second segment were then averaged together to get a single representative PSD for the 30 minute recording session. Total power (dB) per frequency range was calculated using the following ranges: delta (Δ) = 1-4 Hz, theta (θ) = 5-10 Hz, alpha (α) = 11-14 Hz, beta (β) = 15-30 Hz, low gamma (lγ) = 45-65 Hz, and high gamma (hγ) 70-90 Hz (Catanese, Carmichael, & van der Meer, 2016; McCracken & Grace, 2009). To account for the 60 Hz notch filter, power values of frequencies from 59 to 61 Hz were not included in the analysis. The power per frequency band was then normalized as a percent of the average total power of the signal from 1 to 90 Hz (beginning of Δ to end of hγ).

Coherence was computed using the function *mscohere* with a 1.3 second sliding Hamming window with 50% overlap. The average coherence between each pair of frequency bands from 1 to 90 Hz (excluding values corresponding to the 60 Hz notch filter) were used to normalize the average coherence of each frequency band within that neural site pair.

### Brain stimulation

To deliver stimulation, a current-controlled stimulator (*PlexStim, Plexon, Plano, TX*) was used to generate a continuous train of biphasic pulses (90 μsec pulse width, 130 Hz, 200 μA). We chose 130 Hz stimulation because this frequency matches what was used in a clinical study of DBS for alcohol dependence (Voges et al., 2013). The output of the stimulator (current and voltage) was verified visually for each animal before and after each stimulation session using a factory-calibrated oscilloscope (TPS2002C, Tektronix, Beaverton, OR). Stimulation was initiated immediately before animals had access to alcohol and turned off at the completion of the 90-minute stimulation session. Animals were exposed to NAcSh or mPFC brain stimulation for 3 sessions in a counterbalanced fashion, followed by a washout period, such that all animals were exposed to both NAcSh and mPFC stimulation. During the washout periods, animals were again allowed to drink alcohol while plugged in to the stimulation cables but without stimulation turned on. It is important to note that a subset of animals also received stimulation to the NAcSh (n=6) and mPFC (n=7) at 20 Hz with washout periods between the two different stimulation parameters. We piloted the impact of low frequency stimulation on alcohol drinking behavior because the frequency of stimulation more accurately mimics clinical TMS, but we did not observe significant changes in drinking behavior in either stimulation target (supplementary Figs. 2A and 2B). Figure 1 outlines the experimental timeline.

**Figure 1.**
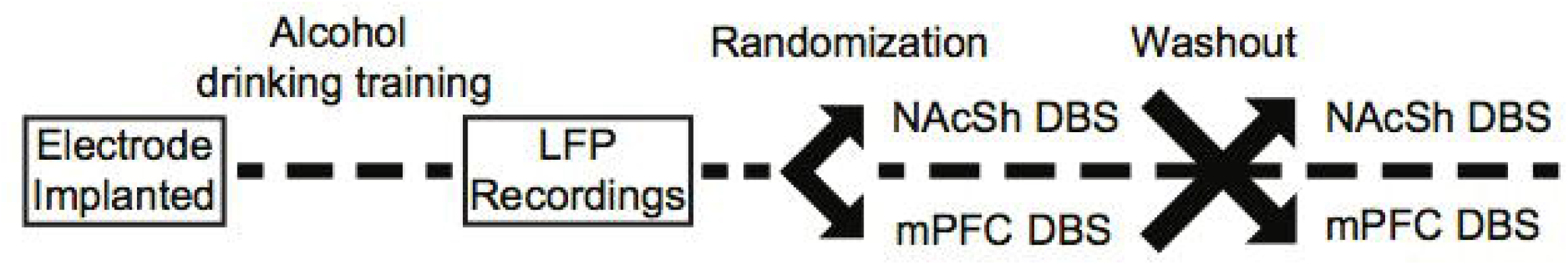
Experimental timeline.

### Statistical Analysis

#### Categorizing rats as HD or LD

The average alcohol consumed (in g/kg) during the last 3 days of the training drinking sessions (prior to stimulation) was calculated for each rat. Rats were subsequently categorized as HD or LD using a median split, which served as the dependent outcome used to build prediction models using the LFP features. A repeated measures ANOVA (RMANOVA) was also used to determine whether levels of alcohol consumption was significantly different between HD and LD across training sessions.

#### Linking corticostriatal activity to HD vs. LD phenotypes

Each recording session produced 60 LFP features: 24 measures of power (6 frequency bands X 4 channels) and 36 measures of coherence (6 frequency bands X 6 channel combinations). We used a penalized regression method (lasso) in order to capture potential combinations of LFP features that correlated with behavioral phenotypes (HD vs. LD). The Matlab package *Glmnet* was used to implement the lasso using a cross-validation scheme with 100 repetitions. The accuracy of the model is reported as the average cross-validated accuracy. We repeated the entire above process on 100 random permutations of the data. The mean accuracy and 95% confidence intervals from the observed and permuted data distributions were determined for comparison.

It is important to note that there are sex differences in alcohol consumption rates in rodents [with females drinking more alcohol than males (Morales, McGinnis, & McCool, 2015)], but the majority of preclinical data has used only male animals (Beery & Zucker, 2011). It is thus unknown whether NAcSh or mPFC circuit oscillations would predict HD vs. LD in females. Since we are aware that the inclusion of females would increase the utility of our models, we also built a model to predict HD vs. LD with 5 extra females included in the median split.

#### Calculating the response to brain stimulation

The overall effect of NAcSh or mPFC stimulation on the amount of alcohol consumed was analyzed using a RMANOVA, with average alcohol consumption (in g/kg) from the 3 days of training and the 3 sessions of stimulation as the within-subjects factors, and group (HD or LD) as the between-subjects factor. Average percent change from training (pre-stimulation) to stimulation was also calculated for each animal and group differences (HD vs. LD) were analyzed with an independent-samples t-test.

### Verification of Electrode Placement

At the end of the experiment, animals were euthanized using CO2 gas and brains were snap frozen in 2-methylbutane on dry ice. Tissue was stored at −20°C prior to sectioning, thionine staining and histologic analysis.

## Results

### Categorization of rats as HD or LD

Using a median split of average alcohol consumed (in g/kg) over the last 3 sessions of limited access to alcohol, 7 rats were categorized as HD and 6 rats were categorized as LD. Figure 2A shows the amount of alcohol consumed during the 12 limited access sessions prior to stimulation stratified by group. A RMANOVA revealed a significant effect of group [F(1,11) = 5.86, p = 0.03, *n^2^p* = 0.35], suggesting that HD and LD drank consistently different amounts of alcohol across training sessions. Figure 2B shows average alcohol consumed during training for the HD and LD, as well as the individual variability in behavior. An independent samples t-test confirmed that average alcohol consumed was significantly different between HD and LD [t(11) = 5.79, p = 0.00].

**Figure 2.**
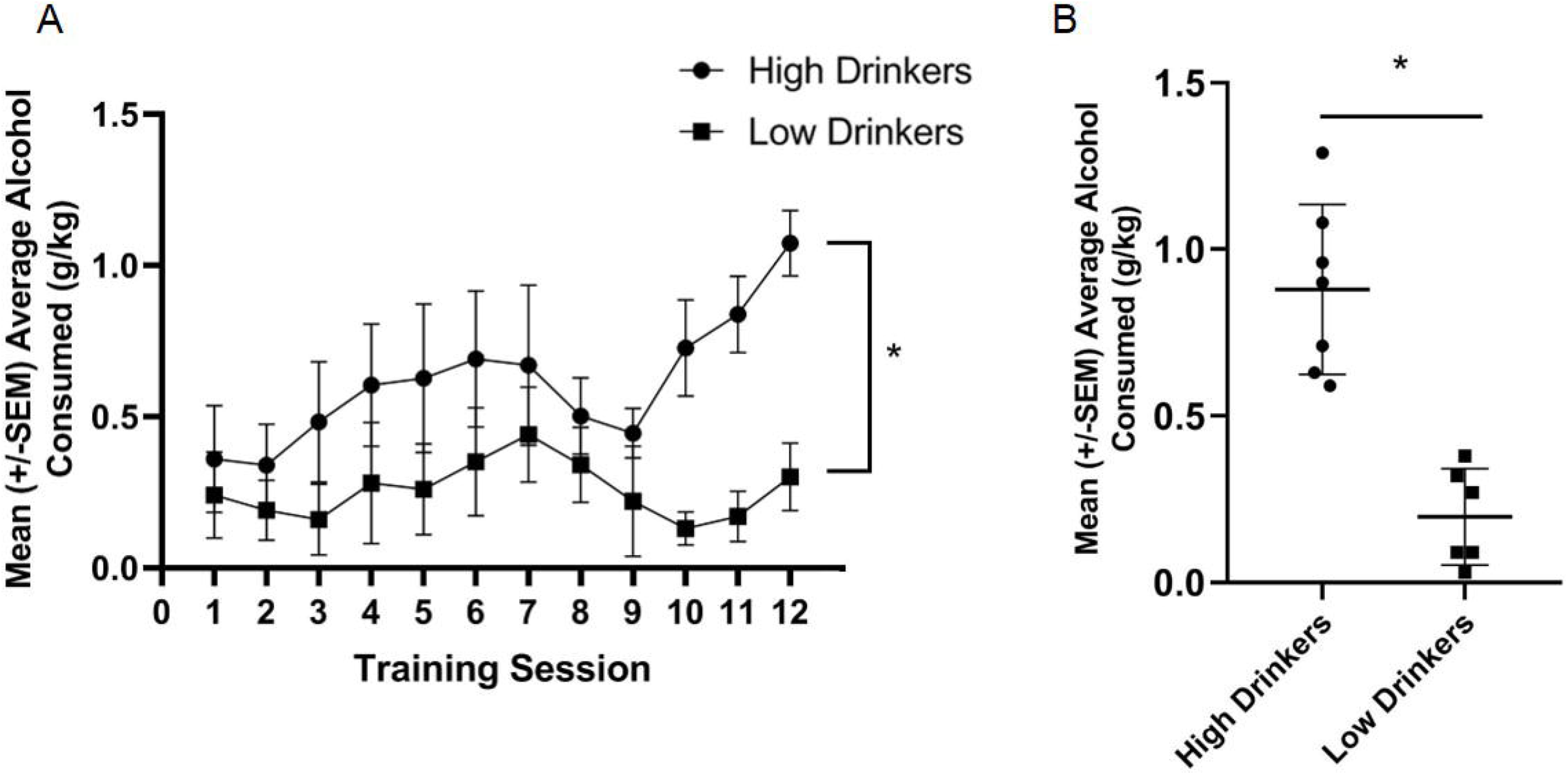
Categorization of rats as HD or LD. Average g/kg of alcohol consumed for HD and LD was significantly different across the 12 training sessions, prior to stimulation [F(1,11) = 5.86, p = 0.03, *n^2^p* = 0.35; **A**]. Average g/kg of alcohol consumed across the last 3 days of alcohol drinking training was significantly different between HD and LD [t(11) = 5.79, p = 0.00; **B**] (n = 6-7/group).

### LFPs recorded within corticostriatal regions predict HD vs. LD

The model built from corticostriatal LFP features was able to outperform permuted data in predicting which rats were HD vs. which rats were LD (permuted μ = 48±1%, real μ = 80±2%; Fig. 3A). Figures 3B and 3C show the precise placement of the electrodes. Interestingly, the top 4/5 neural features important in building the prediction model indicate that gamma power and coherence was high in HD vs. LD (Table 1).

**Figure 3.**
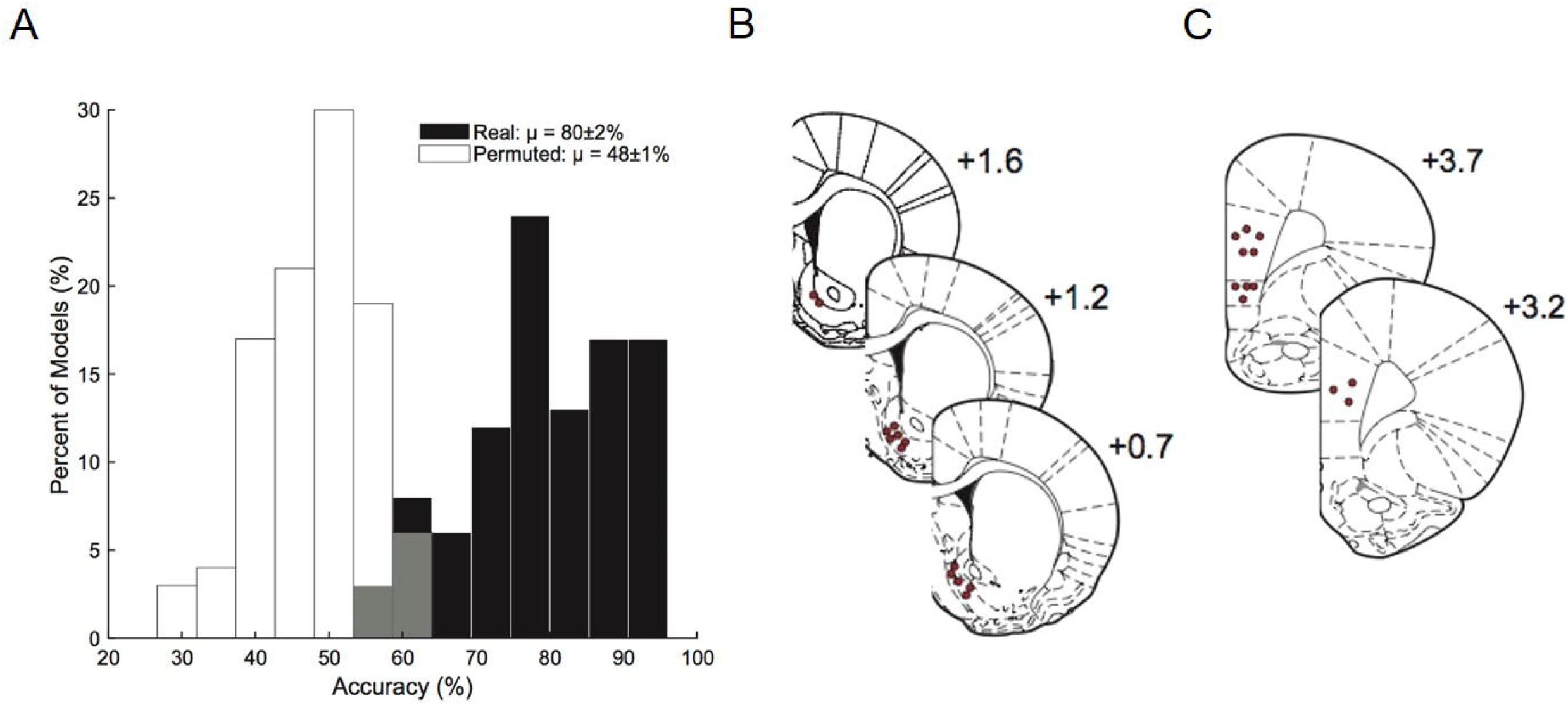
Prediction model. Corticostriatal LFP oscillations predict HD vs. LD better than permuted data (permuted μ = 48±1%; real μ = 80±2%; **A**). Histology figures representing electrode placements in the NAcSh (**B**) and mPFC (**C;** n = 10-14/group).

**Table 1.**
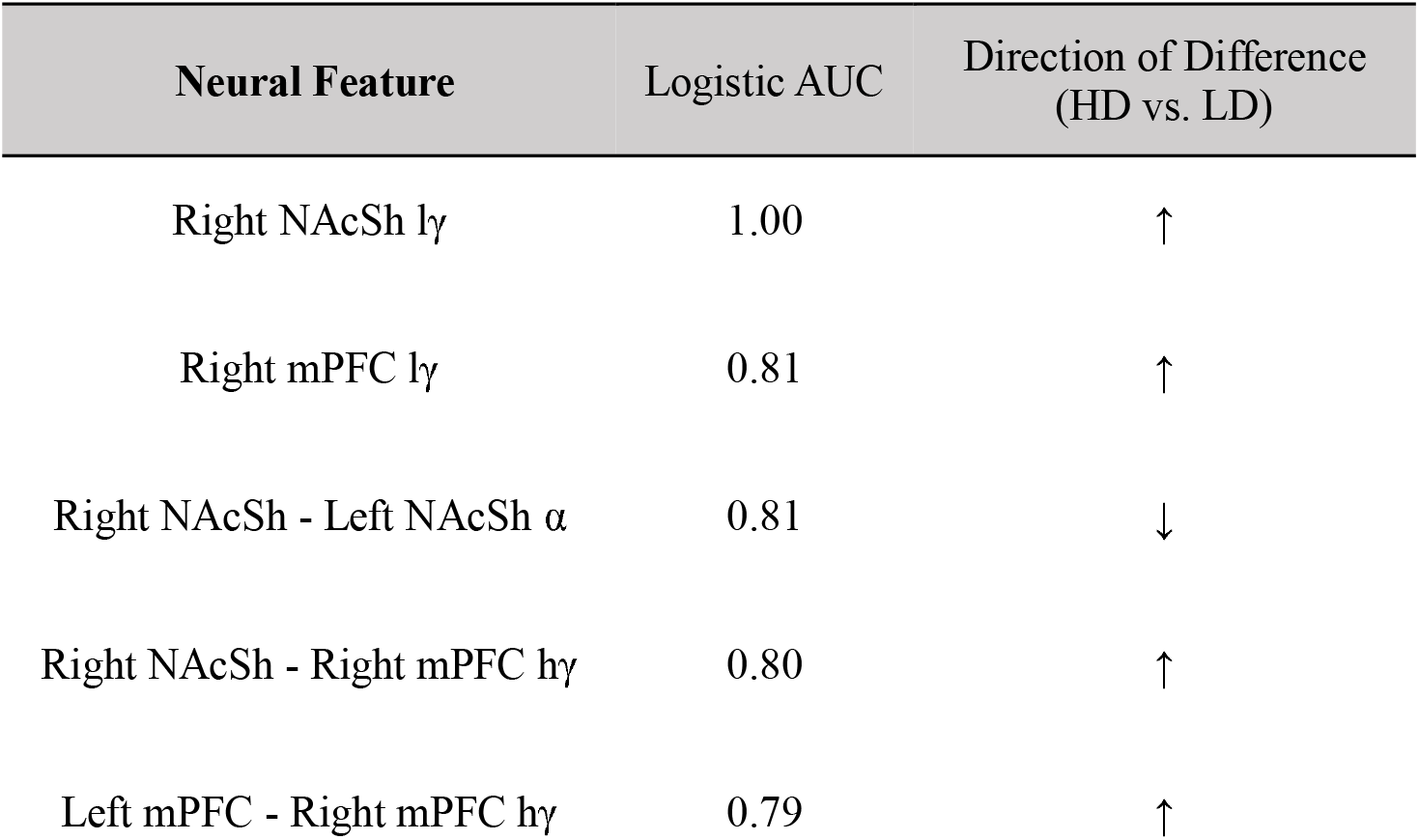
Neural features important in model prediction accuracies. The top five LFP features used in models predicting HD vs. LD. Frequency bands [alpha (α), low gamma (lγ), and high gamma (hγ)] are described for power features within and coherence features between neural sites. Arrows indicate the state of HD relative to LD.

Supplementary Fig. 1 shows the predictive model with 5 extra females included (n = 2 HD and 3 LD). This combined model built from male and female corticostriatal LFP features was able to outperform permuted data in predicting which rats were HD vs. which rats were LD (permuted μ = 47±1%, real μ = 61±2%), but the accuracy significantly drops from the “male-only” model (80% to 61%). This preliminary data suggests there may be sex differences in the neural features that predict HD vs. LD, but future work is necessary to fully test this hypothesis.

### NAcSh stimulation significantly reduces alcohol consumption in HD

A RMANOVA showed a significant time*group interaction [F(1,11) = 5.89, p = 0.03, *n^2^p* = 0.35] for NAcSh stimulation on alcohol consumption (Fig. 4A), which suggests that HD, but not LD, showed a significant decrease in alcohol drinking due to stimulation on a population level. It is important to note, however, that individual response to NAcSh stimulation in HD was largely driven by significant changes in only a couple of rats (Fig. 4B). Individual responses to NAcSh stimulation in LD suggests that this subpopulation was largely unaffected (Fig. 4C).

**Figure 4.**
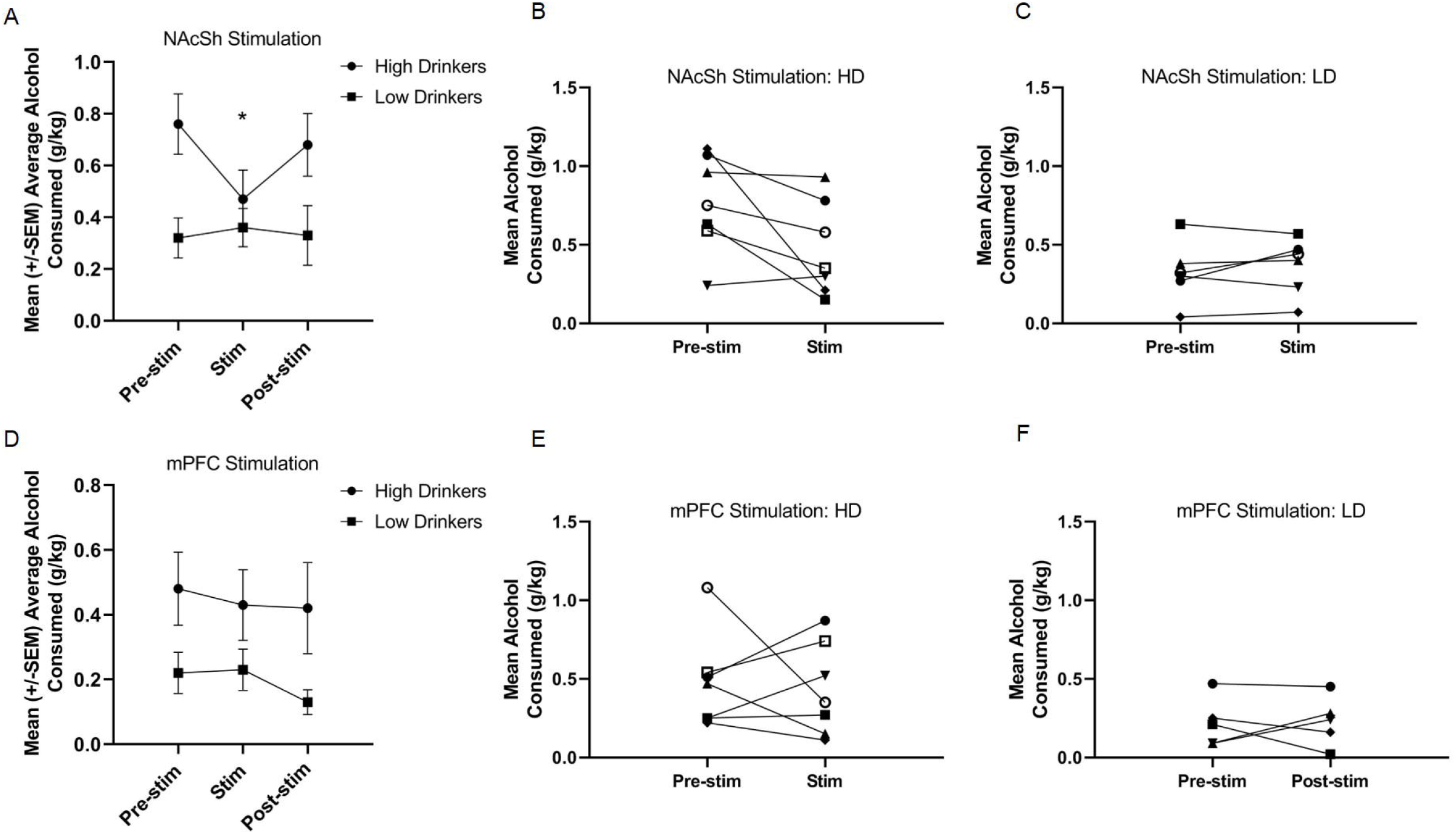
Response to 130 Hz NAcSh and mPFC stimulation. NAcSh stimulation led to a significant decrease in alcohol consumption from training (pre-stimulation) to the stimulation sessions in the HD only [F(1,11) = 5.89, p = 0.03, *n^2^p* = 0.35; **A**]. Panels **B** and **C** represent the individual responses to NAcSh stimulation in HD and LD, respectively. Neither HD or LD showed a significant change in alcohol consumption due to mPFC stimulation [F(1,10) = 0.04, p = 0.85, *n^2^p* = 0.00; **D**]. Panels **E** and **F** represent the individual responses to mPFC stimulation in HD and LD, respectively (n = 5-7/group).

A RMANOVA showed no effect of time [F(1,10) = 0.04, p = 0.85, *n^2^p* = 0.00] or group [F(1,10) = 4.0, p = 0.08, *n^2^p* = 0.28], nor a group*time interaction [F(1,10) = 0.08, p = 0.78, *n^2^p* = 0.01] for mPFC stimulation on alcohol consumption (Fig. 4D). Figs. 4E and 4F show the individual responses to mPFC stimulation in HD and LD, respectively. Due to electrode headcap failure, one animal in the NAc group did not receive 130Hz mPFC stimulation.

## Discussion

Here we show that corticostriatal oscillations can be used to classify rats as HD or LD, indicating that this circuit contains information regarding vulnerability to excessive alcohol consumption. Additionally, when we perturbed the corticostriatal circuit with 130 Hz NAcSh stimulation, only HD rats showed a significant decrease in alcohol consumption. Interestingly, however, a closer look at the individual responses to NAcSh stimulation suggest that the significant population effect is driven by only a couple of rats. These data highlight an important caveat impeding more widespread use of circuit-based therapies in clinical populations: highly variable treatment outcomes across individuals. The fact that many of our rats did not show decreases in alcohol consumption with NAcSh stimulation suggests that perhaps not all individuals respond to stimulation of the same brain target [as we have previously observed in a model of binge eating (Doucette et al., 2018)] or the same stimulation parameters. This caveat does not necessarily diminish the impact of the current results, but indicate that further advancement of circuit-based interventions requires that electrode target selection and stimulation parameters be personalized based on an individual’s unique brain structure and function. Additional research is needed to determine whether the personalization of circuit-based intervention using network activity can lead to consistent and meaningful improvements in stimulation outcomes in preclinical models of other neuropsychiatric conditions.

One interesting outcome of this study is that the majority of neural features important in predicting HD vs. LD were observed within the gamma frequency range (lγ and hγ), where HD rats showed increased γ power and coherence compared to LD rats. Indeed, γ oscillations from the NAc have been previously correlated with reward-related behaviors in rats (Dejean et al., 2017; van der Meer & Redish, 2009), and can be altered by dopamine manipulation (Berke, 2009; Lemaire et al., 2012). The current data suggest that γ oscillations from the NAc may also provide an important readout regarding vulnerability to develop excessive alcohol consumption, but future work is necessary to begin to causally relate these neural signatures with behavioral phenotypes.

Previous studies have also demonstrated that NAc stimulation can significantly reduce alcohol consumption in rodents (Henderson et al., 2010; Knapp et al., 2009). These data, in conjunction with the present experiment, suggest that NAc stimulation can reduce alcohol consumption in a subset of individuals. The NAc is an important nexus point within a complex network including reciprocal projections to frontal brain regions involved in behavioral decisionmaking (Goto & Grace, 2008). It is thus not surprising that the NAc is a commonly proposed target for treating addiction-related behaviors using circuit-based interventions. However, future studies will need to continue to add to a growing literature linking individual variation in systems-level brain activity to variation in treatment outcomes for alcohol use disorders.

To our knowledge, the present experiment is the first to assess the efficacy of mPFC stimulation to modulate alcohol consumption in rats, though it has been investigated to alleviate treatment-resistant depression. Alcohol dependence and depression are highly co-morbid (Grant & Harford, 1995) and the neural circuits underlying both diseases significantly overlap (Pujara & Koenigs, 2014). Importantly, a recent clinical pilot study showed that TMS of the mPFC reduced craving and self-reported alcohol consumption in a group of alcoholic individuals (Ceccanti et al., 2015). While our work does not directly support the therapeutic potential of mPFC stimulation for alcohol consumption, we tested only one of an infinite set of stimulation parameters, and future clinical and preclinical research would need to parse this parameter space to determine the potential of mPFC stimulation targets for treating addictive disorders.

We would like to acknowledge several limitations in this study. First, this study is limited by a small sample size, but used statistical methods (lasso) to shrink our predictor set and through permutation controls have attempted to account for the effects of overfitting. Secondly, we aimed our mPFC electrodes to the infralimbic/prelimbic junction as we did not have an *a priori* hypothesis that one site would be more effective in reducing alcohol drinking over the other. Furthermore, the electrical field of our stimulation [~1 mm radius (Hamani et al., 2014)] may have affected targets outside of the intended neural structures. We did not observe any significant correlations between placement of the electrodes and response to stimulation (see supplemental Table 1), but future research could be designed to more specifically evaluate differences between infralimbic and prelimbic stimulation on alcohol drinking behavior. Third, though others have demonstrated that NAc or mPFC stimulation does not alter locomotor activity in rodents (Guo et al., 2013; Laver, Diwan, Nobrega, & Hamani, 2014), we did not directly measure locomotor behavior, and it may be possible that stimulation altered alcohol consumption through a non-specific mechanism (decreased activity). We also did not measure natural rewards during stimulation, but others have demonstrated that NAc or mPFC stimulation does not affect water or sucrose intake, respectively (Levy et al., 2007; Luigjes et al., 2012), suggesting that any reductions in alcohol drinking behavior we observed were not simply related to reductions in consummatory behavior in general.

Overall, the current study is an example of how systems-level brain activity might be utilized as a tool for identifying vulnerable subpopulations to target for therapeutic interventions. Future treatments for alcohol dependence and other addictive disorders could also use similar electrophysiological and unbiased computational methods to match effective therapies to the appropriate subpopulations to decrease variability in treatment outcomes.

## Acknowledgments

This work was supported by funds from the Department of Psychiatry at the Geisel School of Medicine at Dartmouth (AIG), the Hitchcock Foundation (WTD, AMH), an LRP grant from the NIH NCATS (WTD) and the Dartmouth Clinical and Translational Science Institute, under award number KL2TR001088 from the National Center for Advancing Translational Sciences (NCATS) of the National Institutes of Health (NIH). This work is currently available as a preprint at: *bioRxiv doi: 10.1101/293597*.

## Author Contributions

AH contributed to planning the experimental design, collected the data and wrote the manuscript. LD analyzed the local field potential data and contributed to editing the manuscript. ND, AS, and MRJ assisted with data collection. WD and AG contributed to planning the experimental design and editing the manuscript.

## Figure Legends

**Supplementary Figure 1.** Prediction model with males and females. Corticostriatal LFP oscillations predict HD vs. LD better than permuted data (permuted μ = 47±1%; real μ = 61±2%).

**Supplementary Figure 2.** Response to 20Hz NAcSh and mPFC stimulation. Neither 20Hz NAcSh [F(1,4) = 0.43, p = 0.85, *n^2^p* = 0.01; **A**] nor mPFC [F(1,4) = 0.79, p = 0.43, *n^2^p* = 0.17; **B**] stimulation altered alcohol consumption from training to the stimulation sessions (n = 2-4/group).

### Table Legends

**Supplementary Table 1.** Histology correlations. Correlations coefficients (*r^2^*) and p values for the relationship between the electrode placements (NAcSh AP coordinate or mPFC DV (infralimbic vs. prelimbic) coordinate and average percent change in g/kg of alcohol consumed during NAcSh or mPFC stimulation. There were no significant correlations between the electrode coordinates and the response to stimulation.

|  | NAcSh AP coordinate |  | mPFC DV coordinate<br>(PL or IL) |  |
| --- | --- | --- | --- | --- |
| Average percent change | $r^2$ | $p$ | $r^2$ | $p$ |
| High Drinkers | 0.27 | 0.56 | -0.63 | 0.13 |
| Low Drinkers | -0.36 | 0.48 | 0.27 | 0.66 |

## References

Beery, A. K., & Zucker, I. (2011). Sex bias in neuroscience and biomedical research. Neuroscience and Biobehavioral Reviews, 35(3), 565–572. https://doi.org/10.1016/j.neubiorev.2010.07.002

Berke, J. D. (2009). Fast oscillations in cortical-striatal networks switch frequency following rewarding events and stimulant drugs. The European Journal of Neuroscience, 30(5), 848–859. https://doi.org/10.1111/j.1460-9568.2009.06843.x

Broadwater, M. A., Lee, S.-H., Yu, Y., Zhu, H., Crews, F. T., Robinson, D. L., & Shih, Y.-Y. I. (2018). Adolescent alcohol exposure decreases frontostriatal resting-state functional connectivity in adulthood. Addiction Biology, 23(2), 810–823. https://doi.org/10.1111/adb.12530

Camchong, J., Stenger, A., & Fein, G. (2013). Resting-State Synchrony in Long-Term Abstinent Alcoholics. Alcoholism: Clinical and Experimental Research, 37(1), 75–85. https://doi.org/10.1111/j.1530-0277.2012.01859.x

Catanese, J., Carmichael, J. E., & van der Meer, M. A. A. (2016). Low- and high-gamma oscillations deviate in opposite directions from zero-phase synchrony in the limbic corticostriatal loop. Journal of Neurophysiology, 116(1), 5–17. https://doi.org/10.1152/jn.00914.2015

Ceccanti, M., Inghilleri, M., Attilia, M. L., Raccah, R., Fiore, M., Zangen, A., & Ceccanti, M. (2015). Deep TMS on alcoholics: effects on cortisolemia and dopamine pathway modulation. A pilot study. Canadian Journal of Physiology and Pharmacology, 93(4), 283–290. https://doi.org/10.1139/cjpp-2014-0188

Centers for Disease Control and Prevention. (2013). Alcohol Related Disease Impact (ARDI) application. Retrieved October 29, 2018, from https://nccd.cdc.gov/DPH_ARDI/Default/Report.aspx?T=AAM&P=f6d7eda7-036e-4553-9968-9b17ffad620e&R=d7a9b303-48e9-4440-bf47-070a4827e1fd&M=8E1C5233-5640-4EE8-9247-1ECA7DA325B9&F=&D=

Costanzo, P. R., Malone, P. S., Belsky, D., Kertesz, S., Pletcher, M., & Sloan, F. A. (2007). Longitudinal differences in alcohol use in early adulthood. Journal of Studies on Alcohol and Drugs, 68(5), 727–737. Retrieved from http://www.ncbi.nlm.nih.gov/pubmed/17690807

Dejean, C., Sitko, M., Girardeau, P., Bennabi, A., Caillé, S., Cador, M., … Le Moine, C. (2017). Memories of Opiate Withdrawal Emotional States Correlate with Specific Gamma Oscillations in the Nucleus Accumbens. Neuropsychopharmacology, 42(5), 1157–1168. https://doi.org/10.1038/npp.2016.272

Doucette, W. T., Dwiel, L., Boyce, J. E., Simon, A. A., Khokhar, J. Y., & Green, A. I. (2018). Machine Learning Based Classification of Deep Brain Stimulation Outcomes in a Rat Model of Binge Eating Using Ventral Striatal Oscillations. Frontiers in Psychiatry, 9, 336. https://doi.org/10.3389/fpsyt.2018.00336

Doucette, W. T., Khokhar, J. Y., & Green, A. I. (2015). Nucleus accumbens deep brain stimulation in a rat model of binge eating. Translational Psychiatry, 5(12), e695. https://doi.org/10.1038/tp.2015.197

Goldstein, R. Z., & Volkow, N. D. (2002). Drug addiction and its underlying neurobiological basis: Neuroimaging evidence for the involvement of the frontal cortex. American Journal of Psychiatry, 159(10), 1642–1652. https://doi.org/10.1038/jid.2014.371

Goto, Y., & Grace, A. A. (2008). Limbic and cortical information processing in the nucleus accumbens. Trends in Neurosciences, 31(11), 552–558. https://doi.org/10.1016/j.tins.2008.08.002

Grant, B. F., & Harford, T. C. (1995). Comorbidity between DSM-IV alcohol use disorders and major depression: results of a national survey. Drug and Alcohol Dependence, 39, 197–206. https://doi.org/10.1016/0376-8716(95)01160-4

Guo, L., Zhou, H., Wang, R., Xu, J., Zhou, W., Zhang, F., … Jiang, J. (2013). DBS of nucleus accumbens on heroin seeking behaviors in self-administering rats. Drug and Alcohol Dependence, 129(1–2), 70–81. https://doi.org/10.1016/j.drugalcdep.2012.09.012

Hägele, C., Friedel, E., Kienast, T., & Kiefer, F. (2014). How Do We “Learn” Addiction? Risk Factors and Mechanisms Getting Addicted to Alcohol. Neuropsychobiology, 70(2), 67–76. https://doi.org/10.1159/000364825

Hamani, C., Amorim, B. O., Wheeler, A. L., Diwan, M., Driesslein, K., Covolan, L., … Nobrega, J. N. (2014). Deep brain stimulation in rats: different targets induce similar antidepressant-like effects but influence different circuits. Neurobiology of Disease, 71, 205–214. https://doi.org/10.1016/j.nbd.2014.08.007

Henderson, M. B., Green, A. I., Bradford, P. S., Chau, D. T., Roberts, D. W., & Leiter, J. C. (2010). Deep brain stimulation of the nucleus accumbens reduces alcohol intake in alcohol-preferring rats. Neurosurgical Focus, 29(2), E12. https://doi.org/10.3171/2010.4.F0CUS10105

Knapp, C. M., Tozier, L., Pak, A., Ciraulo, D. A., & Kornetsky, C. (2009). Deep brain stimulation of the nucleus accumbens reduces ethanol consumption in rats. Pharmacology, Biochemistry, and Behavior, 92(3), 474–479. https://doi.org/10.1016/j.pbb.2009.01.017

Koob, G. F., & Volkow, N. D. (2010). Neurocircuitry of addiction. Neuropsychopharmacology: Official Publication of the American College of Neuropsychopharmacology, 35(1), 217–238. https://doi.org/10.1038/npp.2010.4

Laver, B., Diwan, M., Nobrega, J. N., & Hamani, C. (2014). Augmentative therapies do not potentiate the antidepressant-like effects of deep brain stimulation in rats. Journal of Affective Disorders, 161, 87–90. https://doi.org/10.1016/j.jad.2014.03.007

Lemaire, N., Hernandez, L. F., Hu, D., Kubota, Y., Howe, M. W., & Graybiel, A. M. (2012). Effects of dopamine depletion on LFP oscillations in striatum are task- and learning-dependent and selectively reversed by L-DOPA. Proceedings of the National Academy of Sciences of the United States of America, 109(44), 18126–18131. https://doi.org/10.1073/pnas.1216403109

Levy, D., Shabat-Simon, M., Shalev, U., Barnea-Ygael, N., Cooper, A., & Zangen, A. (2007). Repeated electrical stimulation of reward-related brain regions affects cocaine but not “natural” reinforcement. The Journal of Neuroscience□: The Official Journal of the Society for Neuroscience, 27(51), 14179–14189. https://doi.org/10.1523/JNEUROSCI.4477-07.2007

Luigjes, J., van den Brink, W., Feenstra, M., van den Munckhof, P., Schuurman, P. R., Schippers, R., … Denys, D. (2012). Deep brain stimulation in addiction: a review of potential brain targets. Molecular Psychiatry, 17(6), 572–583. https://doi.org/10.1038/mp.2011.114

Matošić, A., Marušić, S., Vidrih, B., Kovak-Mufić, A., & Cicin-Šain, L. (2016). NEUROBIOLOGICAL BASES OF ALCOHOL ADDICTION. Acta Clinica Croatica, 55(1), 134–150. Retrieved from http://www.ncbi.nlm.nih.gov/pubmed/27333729

McCracken, C. B., & Grace, A. A. (2009). Nucleus Accumbens Deep Brain Stimulation Produces Region-Specific Alterations in Local Field Potential Oscillations and Evoked Responses In Vivo. Journal of Neuroscience, 29(16), 5354–5363. https://doi.org/10.1523/JNEUROSCI.0131-09.2009

Morales, M., McGinnis, M. M., & McCool, B. A. (2015). Chronic ethanol exposure increases voluntary home cage intake in adult male, but not female, Long-Evans rats. Pharmacology, Biochemistry, and Behavior, 139(Pt A), 67–76. https://doi.org/10.1016/j.pbb.2015.10.016

Morganstern, I., Liang, S., Ye, Z., Karatayev, O., & Leibowitz, S. F. (2012). Disturbances in behavior and cortical enkephalin gene expression during the anticipation of ethanol in rats characterized as high drinkers. Alcohol, 46(6), 559–568. https://doi.org/10.1016/j.alcohol.2012.05.003

Pratt, L. M., Gates, S. K. E., Smith, B. R., & Amit, Z. (2002). A relation between maze performance and increased ethanol intake in Long-Evans rats. Alcohol (Fayetteville, N.Y.), 26(2), 121–126. Retrieved from http://www.ncbi.nlm.nih.gov/pubmed/12007587

Pujara, M., & Koenigs, M. (2014). Mechanisms of reward circuit dysfunction in psychiatric illness: prefrontal-striatal interactions. The Neuroscientist□: A Review Journal Bringing Neurobiology, Neurology and Psychiatry, 20(1), 82–95. https://doi.org/10.1177/1073858413499407

Sharko, A. C., Kaigler, K. F., Fadel, J. R., & Wilson, M. A. (2013). Individual Differences in Voluntary Ethanol Consumption Lead to Differential Activation of the Central Amygdala in Rats: Relationship to the Anxiolytic and Stimulant Effects of Low Dose Ethanol. Alcoholism: Clinical and Experimental Research, 37, E172–E180. https://doi.org/10.1111/j.1530-0277.2012.01907.x

Spoelder, M., Flores Dourojeanni, J. P., de Git, K. C. G., Baars, A. M., Lesscher, H. M. B., & Vanderschuren, L. J. M. J. (2017). Individual differences in voluntary alcohol intake in rats: relationship with impulsivity, decision making and Pavlovian conditioned approach. Psychopharmacology, 234(14), 2177–2196. https://doi.org/10.1007/s00213-017-4617-6

van der Meer, M. A. A., & Redish, A. D. (2009). Low and high gamma oscillations in rat ventral striatum have distinct relationships to behavior, reward, and spiking activity on a learned spatial decision task. Frontiers in Integrative Neuroscience, 3, 9. https://doi.org/10.3389/neuro.07.009.2009

Voges, J., Müller, U., Bogerts, B., Münte, T., & Heinze, H.-J. (2013). Deep Brain Stimulation Surgery for Alcohol Addiction. World Neurosurgery, 80(3–4), S28.e21–S28.e31. https://doi.org/10.1016/j.wneu.2012.07.011

Wilhelm, C. J., & Mitchell, S. H. (2008). Rats bred for high alcohol drinking are more sensitive to delayed and probabilistic outcomes. Genes, Brain and Behavior, 7(7), 705–713. https://doi.org/10.1111/j.1601-183X.2008.00406.x

