## Supplementary figures and images for "Corticostriatal oscillations predict high vs. low drinkers in a preclinical model of limited access alcohol consumption"

### Supplemental Fig 1

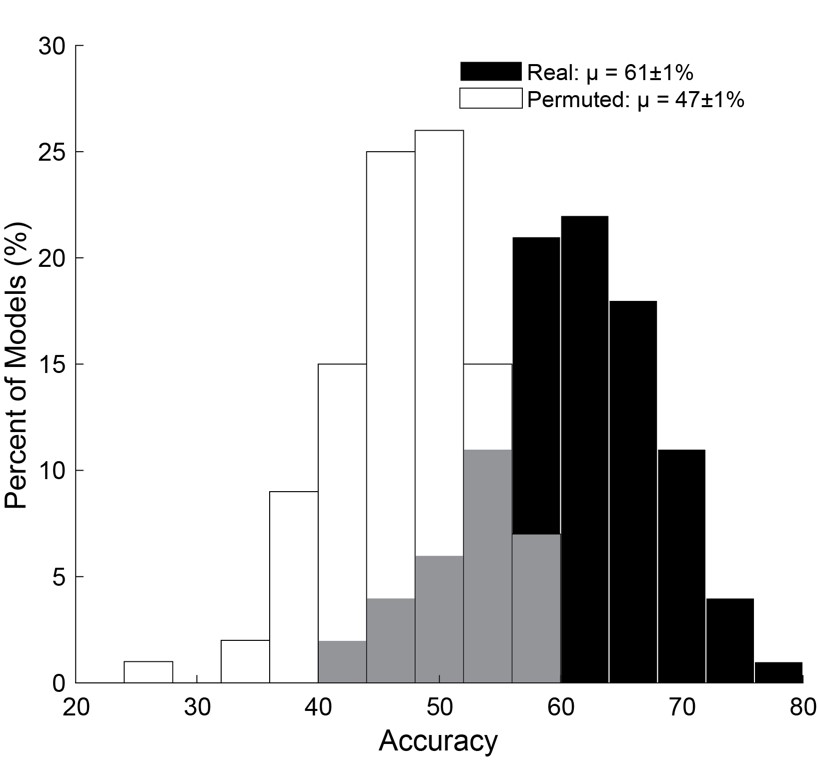

### Supplemental Figure 2

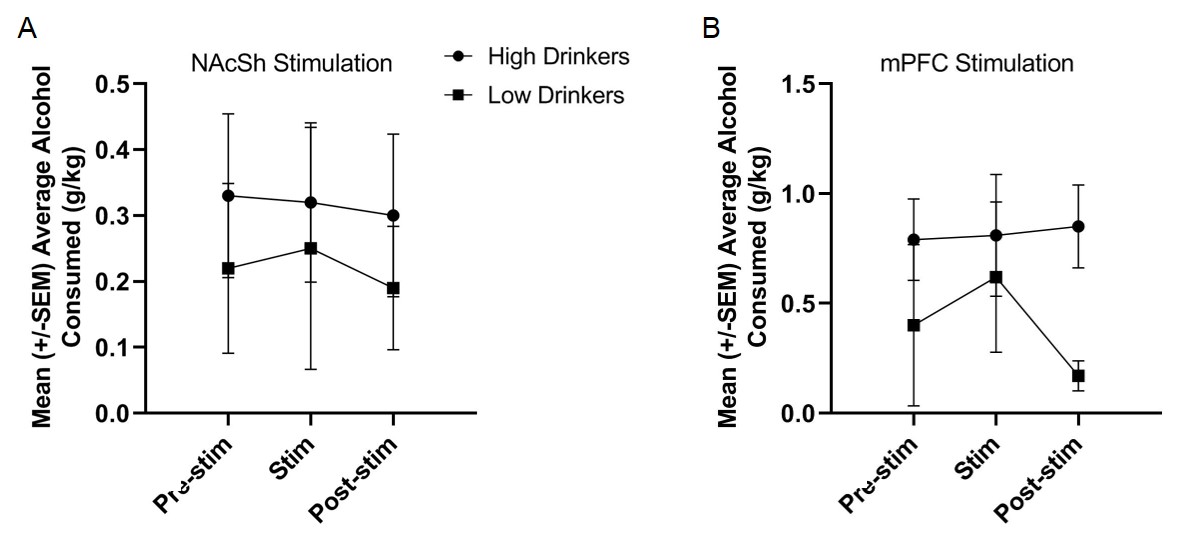
